# A Priori Activation of Apoptosis Pathways of Tumor (AAAPT) Technology: Development of Targeted Apoptosis Initiators for Cancer Treatment

**DOI:** 10.1101/844605

**Authors:** Raghu Pandurangi, Marco Tomasetti, Thillai Verapazham Sekar, Ramasamy Paulmurugan, Cynthia Ma, Sandeep, Manjushree Anjanappa, Harikrishna Nakshatri

**Affiliations:** Sci-Engi-Medco Solutions Inc., 573, Lexington Landing Pl, St Charles, MO 63303-1750; Department of Clinical and Molecular Sciences, Section of Experimental and Occupational Medicine, Polytechnic University of Marche, Via Tronto 10/A, 60126, Ancona, Italy; Molecular Imaging Program at Stanford (MIPS), Canary Center for Cancer Early Detection, Department of Radiology, Stanford University School of Medicine, 3155 Porter Drive, Room: 2236, Palo Alto, CA 94304; Siteman Cancer Center, Washington University School of Medicine, St. Louis, MO 63110, USA

## Abstract

Cancer cells develop tactics to circumvent the interventions by desensitizing themselves to interventions. The principle routes of desensitization include a) activation of survival pathways (e.g. NF-kB, PARP) and b) downregulation of cell death pathways (e.g. CD95/CD95L). As a result, it requires high therapeutic dose to achieve tumor regression which, in turn damages normal cells through the collateral damaging effects. Methods are needed to sensitize the low and non-responsive resistant tumor cells including cancer stem cells (CSCs) in order to evoke a better response from the current treatments. Current treatments including chemotherapy can induce cell death only in bulk cancer cells sparing CSCs and cancer resistant cells (CRCs) which are shown to be responsible for high recurrence of disease and low patient survival. Here, we report several novel tumor targeted sensitizers derived from the natural Vitamin E analogue (AMP-001-003). The drug design is based on a novel concept “A priori activation of apoptosis pathways of tumor technology (AAAPT) which is designed to activate specific cell death pathways and inhibit survival pathways simultaneously. Our results indicate that AMP-001-003 sensitize various types of cancer cells including MDA-MB-231 (triple negative breast cancer), PC3 (prostate cancer) and A543 (ling cancer) cells resulting in reducing the IC-50 of doxorubicin *in vitro*. At higher dose, AMP-001 acts as an anti-tumor agent on its own. The synergy between AMP-001 and doxorubicin could pave a new pathway to use AMP-001 as a neoadjuvant to chemotherapy to achieve a better efficacy and reduced off-target toxicity by the current treatments.

**Summary Statement:** A Priori Activation of Apoptosis Pathways of Tumor often referred to as “AAAPT” is a novel targeted tumor sensitizing technology which synergizes with chemotherapy to enhance the treatment efficacy.

## Introduction

Tumor cells have a remarkable ability to circumvent endogenous and exogenous toxicities by deactivating cell death pathways and thereby desensitizing themselves to interventions^1^. Defects in apoptosis pathways (e.g. CD95, APO-1/Fas) would make tumor cells insensitive to chemotherapy^2^. For example, loss of CD95 signaling pathway resulted in the development of cancer^3^ and resistance to chemotherapy^4^. Restoration of the apoptotic machinery with apoptosis inducing ligands (apoptogens) is an area of active investigation in the drug development. Several agents have been developed to activate TRAIL^5^, to downregulate Bcl-2^6^ and to restore the function of the mutated p53 in order to sensitize tumor cells. Similarly, new advances in the molecular biology of cancer cells reveal that cancer stem cells (CSCs) and cancer resistant cells (CRCs) escape cell death by activating the survival pathway (e.g. NF-kB) and by hyperacting DNA repair enzyme Poly ADP ribose polymerase (PARP) which repairs DNA breaks caused by oncology drugs. Tumor sensitizing technology, specifically for CSCs and CRCs may have to consider offsetting these pathways. A better response from the existing treatments means reducing the therapeutic dose without compromising efficacy and thus, reduce the dose related off-target toxicity. Consequently, tumor sensitizers, potentially can be used as neoadjuvant to chemotherapy to expand the therapeutic index of chemotherapy^7^.

Tumor sensitivity to chemotherapy *in vivo* is shown to be dependent on spontaneous baseline tumor apoptosis index^8^. Aggressive or incurable cancers show low tumor apoptosis index (AI) and low sensitivity to chemotherapy compared to benign cancer^9^. In fact, the spontaneous levels of apoptosis are strong predictor of treatment response^10^. For example, low baseline apoptosis index (AI) of patient tumors (see Fig 4 in ref 9), is directly correlated to non-respondent patients to chemotherapy ((> 67%). Lower the baseline apoptosis index of tumor, least the response from chemotherapy and vice versa (Fig 3 in ref 2). The overall 5-year survival rates for the patient group with high AI (> 0.97) and low AI (< 0.97, p =0.001) were 89.6 % and 69.2 % respectively^11^ indicating that AI could be a potential biomarker of risk stratification of tumors/patients to see who responds better to which treatments if AAAPT technology goes to clinical level.

We have developed a novel technology, “A priori Activation of Apoptosis Pathways of Tumor” (AAAPT) as a strategy to sensitize low responsive tumor cells in order to evoke a better response from chemotherapy^12^. The goal is to understand the molecular biology of desensitization tactics of tumor cells which bypass the intervention by reactivating specific dysregulated cell death pathways and inhibiting survival pathways simultaneously and selectively in tumor cells sparing normal cells. Since ubiquitous or systemic activation of apoptosis can induce undesirable neurodegeneration and myelosupression^13^, targeting is essential^14^.

Triple negative breast cancer (TNBC) is an aggressive form of breast cancer for which no targeted therapy is approved so far due to lack of biomarkers (ER, PR and HER2 negative) based on which drugs are designed^15^. About 90 % of TNBC patients experience metastatic recurrence within 3-5 years with a median survival of just 13 months^16^. Most front-line treatments for TNBC patients including anthracycline drugs despite they induce cardiotoxicity. Loss of cardiomyocytes cannot be replaced since they do not replicate easily. Hence, it is an unmet medical need to improve the clinical performance of current treatments that lowers the off-target toxicity including cardiotoxicity using new tumor sensitizing technology, particularly for TNBC patients who have limited options of treatment, except chemotherapy. Thus, adjuvant or perioperative chemotherapy and molecule-targeted chemotherapy have been in clinical trials for controlling metastasis, reducing cancer recurrence and increasing overall survival^16b^. TNBC cells are also known to downregulate CD95 pathway which makes them insensitive to chemotherapy^17^. Hence, we investigate the tumor sensitizing potential of leading AAAPT molecules AMP-001/003 in TNBC cells to evoke a better response or synergistic effect with a standard chemotherapy for example, doxorubicin. Doxorubicin (dox) is a first-line of treatment which is widely used in clinical therapeutic regimens for a variety of cancers including TNBC. Nevertheless, the clinical application is limited due to the drug resistance and adverse effects, including cardiomyopathy, typhlitis, and acute myelotoxicity^18^. To decrease dox-induced dose-dependent toxicity, enhancing the effective therapeutic index via combination with nontoxic or selective apoptotic inducing ligands has been investigated. Here, we propose novel targeted tumor sensitizers which address how the cancer cells circumvent interventions by reactivating cell death pathways and inhibiting survival pathways simultaneously. Since desensitization tactics by cancer cells are successful in thwarting the efficacy of treatments, a fundamental reversal of cancer cells tactics may have high impact on several modes of interventions (e.g. chemotherapy, radiation and immunotherapy). Here, we report to demonstrate the synergistic effect of AMP-001-003 with front line chemotherapeutics *in vitro* and *in vivo* in a TNBC animal model.

### Rational Drug Design

The drug design AMP-001 involves pegylation of α-tocopheryl succinate (apoptogen) with a dipeptide linker valine-citrulline (VC) which is cleavable by tumor specific Cathepsin B enzyme giving rise to a biologically active final molecule (Scheme 1). We have also synthesized VC conjugated α-tocopherol as a pro-drog with ether linkage (AMP-002) or ester linkage (AMP-003). AMP-002 and AMP-003 are designed without pegylation, yet cleavable by tumor specific enzyme Cathepsin B. This design provided impetus for a) targeting apoptogen to cancer cells using tumor specific biomarker Cathepsin B^19^ sparing normal cells, b) releasing the drug near tumor sites through the release of the biological component by cleaving at valine-citrulline link, c) pegylation to make it water soluble, d) enhancing the bioavailability of AMP-001and e) to keep it intact as a pro-drug in blood circulation for a long time to reach tumor sites. The rationale behind using cathepsin B cleavable linkers is based on the better safety profile of Cathepsin B cleavable prodrugs doxorubicin^20a,^ paclitaxel^20b^ and antibodies^20c^ (Seattle Genetics) compared to unmodified drugs. Cathepsin B is known to a) be a tumor specific biomarker and b) cleave valine-citrulline substrate^21^. For example, high delineation of tumor from the surrounding tissues by cathepsin B sensitive optical probes^22^ shows that this enzyme is restricted to invading cancers.

**Scheme 1:**
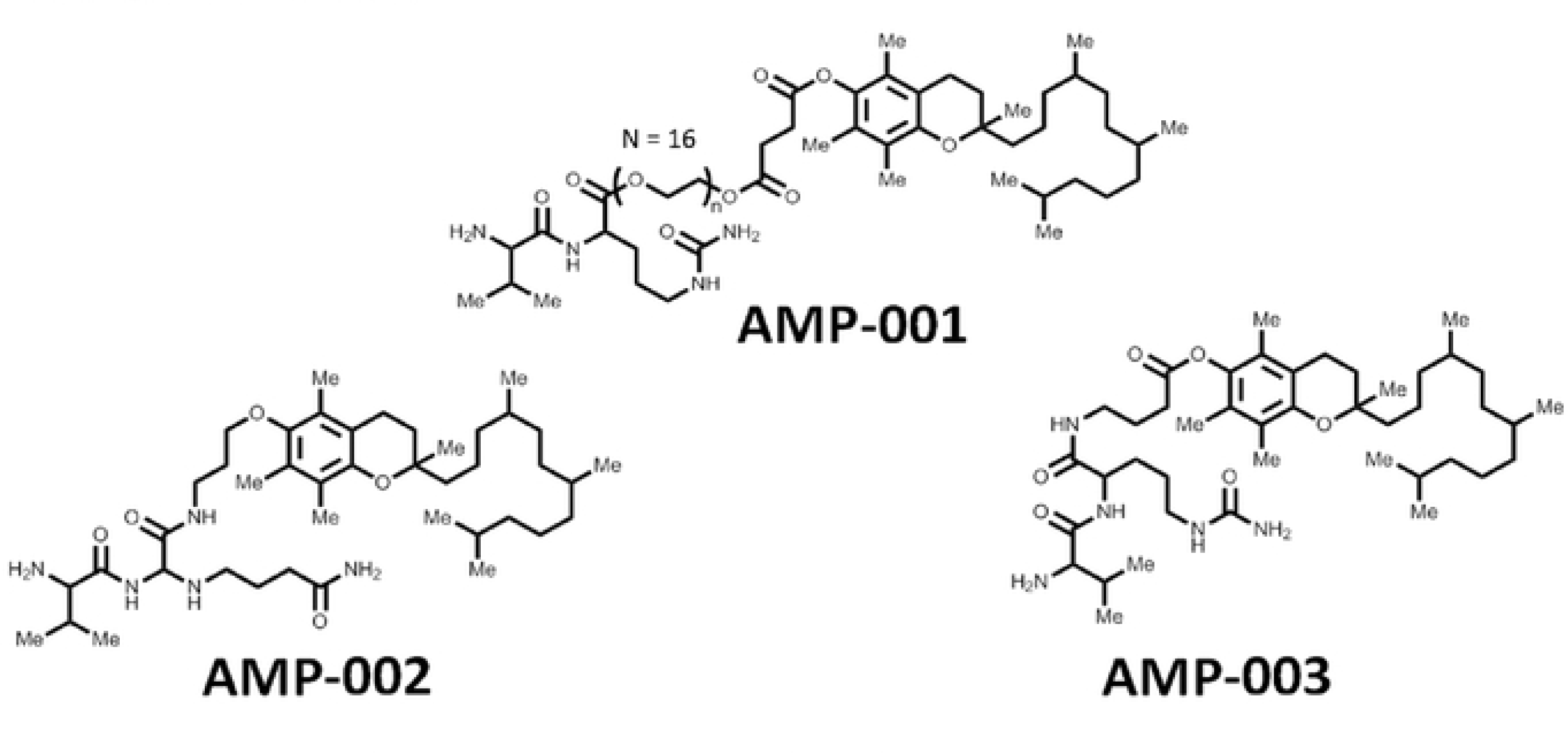
Structures of AMP-001, AMP-002 and AMP-003.

Although, parent molecule α-tocopheryl succinate (α-TOS) showed promising anti-tumor properties *in vitro*, studies in an immunocompetent mouse *in vivo* model showed that α-TOS was not only ineffective at the published doses, but resulted in severe side effects due to lack of targeting^23^. Elevated ROS levels have to be achieved selectively in cancer cells which can render them sensitive to agents that cause oxidative stress^24–25^. There are several chemotherapeutics^26^ which may augment ROS without selectivity leading to non-specific side effects. Hence, targeting is essential.

Here, we examined the effect of combining AMP-001 and doxorubicin for a potential synergistic effect in TNBC both *in vitro* and *in vivo*. These results indicate that low dose pretreatment of tumor *in vivo* prepares tumor to make it sensitive to doxorubicin by showing a cumulative and significant tumor regression compared to individual drugs. A potential mechanism of action is being suggested based on the limited data.

## Materials and Methods

### Cell Culture and Treatment

The TNBC cell lines MDA-MB-231 and MCF-7 were cultured in Dulbecco’s modified Eagle’s medium (DMEM) supplemented with 10% FBS, 100 IU/ml penicillin, 100 μg/ml streptomycin in a humidified atmosphere with 5% CO_2_ at 37℃.

#### NF-kB Inhibition Assay

MDA-MB-231 TNBC cells (5×10^-5^ cells/well) were seeded in 6-well plate and allowed to attach overnight, and treated with tocopheryl succinate (50 µM) or AMP-001/002/003 (20 µM) for 24 h. Alternatively, cells were incubated with the agents (4 h) and NF-kB pathway was activated by TNF-alpha (50 ng/ml) for 30 min. After treatment, cells were harvested, and the pellet re-suspended in the cytoplasm extract (CE) buffer (10 mM HEPES, 60 mM KCl, 1 mM EDTA, 0.075% (v/v) NP40, 1mM DTT and 1 mM PMSF, adjusted to pH 7.6). After 5 min incubation on ice, the pellet was centrifuged at 1500 rpms for 5 min, and the supernatant containing the cytosolic fraction was collected. The remaining pellet was washed in CE buffer, and lysed in the nuclear extract (NE) buffer (20 mM Tris-HCl, 420 mM NaCl, 1.5 mM MgCl2, 0.2 mM EDTA, 1 mM PMSF and 25% (v/v) glycerol, adjusted to pH 8.0). After 20 min incubation on ice, the pellet was centrifuged at 12000 rpms for 10 min at 4°C, and the supernatant was collected. The cytoplasm and nuclear extracts were stored at −80°C until used.

#### Western Blotting

For western blot analysis, protein samples (50 μg per lane) were resolved using 12.5% SDS-PAGE, and transferred to nitrocellulose membranes, and incubated overnight with anti-p65 (Cell Signaling Technology, Danvers, MA). β-Actin and lamin (Bethyl, Montgomery, TX, USA) were used as loading controls for the cytosolic and nuclear fractions, respectively. After incubation with an HRP-conjugated secondary IgG (Sigma), the blots were developed using the ECL detection system (Pierce Biotechnology, Rockford, IL, USA). Band intensities were visualized by ChemiDoc using the Quantity One software (BioRad, Hercules, CA).

### Reagents

The novel compound AMP-001-003 were designed and synthesized by Sci-Engi-Medco Solutions Inc (SEMCO). The drug doxorubicin was purchased from Sigma-Aldrich Biotechnology. For all AMP-001-003/DOX combinational treatment, cancer cells were pretreated with AMP-001-003 before doxorubicin was added into the culture. All above micromolecular reagents were dissolved and saved according to manufacturer specifications.

### Cell Culture

MDA-MB-231 cells (ATCC HTB-26) were cultured in RPMI 1640 culture medium with L-glutamine (Thermo-Fisher Scientific, Waltham MA, Catalog #11875-093), supplemented with 10% fetal bovine serum and 1% Penicillin/Streptomycin (P/S) solution. Cells were incubated in a humidified incubator with 5%CO_2_. MCF10A (CRL-10317) cells were cultured in DMEM /F12 Ham’s Mixture supplemented with 5% Equine Serum (Thermofisher Catalog # 16050130), EGF 20 ng/ml (Sigma), insulin 10μg/ml (Sigma), hydrocortisone 0.5 mg/ml (Sigma), cholera toxin 100 ng/ml (Sigma), 100 units/ml penicillin and 100 μg/ml streptomycin. The cancer cell line A-549 and PC3 were purchased from the ATCC and grown in the RPMI-1640 medium supplemented with 2 mM L-glutamine, 100 U/ml penicillin, 100 µg/ml streptomycin and 10% FBS, at 37°C and 5% CO_2_ in humidified atmosphere. α-TOS (from Sigma) and AMP compounds were dissolved in water (AMP-001), ethanol (AMP-002) and DMSO (AMP-003) respectively, diluted in the complete medium to the final concentration of 60 µM and 30 µM, respectively, and added to cells at 0.1% of the solvent.

### Cell Viability Assay

Cell viability was evaluated by the MTT method. A-549 and PC3 cells were plated in 96-well flat-bottom plates at the density of 2×10^4^ per well, allowed to attach overnight, and treated with α-TOS (60 µM) or the AMP compounds (30 µM) in the presence or absence of NAC and Tiron (10 mM), z-VAD-fmk (10 µM), 3-ABA (2 mM), 3-MA (0.5 mM) or CQ (10 µM). After 24 h, 10 µl (3–4,5-dimethylthiazol-2-yl)-2,5-diphenyltetrazolium bromide (MTT; Sigma) were added to each well and the plate incubated at 37°C for 3 h. After removing the media, 200 µl of isopropanol were added to dissolve the crystals. Absorbance was read at 550 nm in an ELISA plate reader (Sunrise, Tecan, Milan, Italy), and the results expressed as relative change with respect to the controls set as 100%.

### Combination Index Calculations

Combination Indices were calculated using CompuSyn software, V. 1.0 (Biosoft, Cambridge, UK). Drug interactions were classified by determining a combination index (CI) recognized as the standard measure of combination effect based on the Chou-Talalay method. The CI values were obtained over a range of fractional cell kill levels (Fa) from 0.05 to 0.95 (5-95% cell kill). Based on the Chou-Talalay method (1,2,3,4), CI < 1 means synergism, CI = 1 means additivity, and CI > 1 is interpreted as antagonism.

### Assessment of ROS Generation

Intracellular hydrogen peroxide was estimated using the fluorescent dye 2′7′-dichlorofluorescein diacetate (20 µM DCFDA; oxidized by hydrogen peroxide to DCF). A-549 and PC3 cells (10^5^) were seeded in 24-well plates, treated with the drugs as above, and ROS formation was over time detected by a fluorescence plate reader (Infinite F200 PRO, Sunrise, Tecan, Männedorf, Swiss).

### Assessment of Cathepsin B Activity

Media and cells were harvested and processed as described previously^12^. Cathepsin B was measured with the fluorogenic substrate Na-CBZ-L-Arg-L-Arg-7-amido-4-methylcoumarin hydrochloride (Sigma) after a 30-min pretreatment with DTT and EDTA. The predominant form of cathepsin B secreted by tumor cells is the inactive proenzyme, and conditioned medium that was freed of cells contained no detectable, active cathepsin B. Protein concentrations in cell lysates were determined by the Bradford method using the Bio-Rad protein assay dye reagent (Bio-Rad Laboratories, Hercules, CA) according to the instructions of the supplier.

### Fluorescent Microscopy

A-549 and PC3 cells were placed in 6-well plates at 3×10^5^ per well on glass coverslip. The cells were allowed to attach overnight and then incubated 24 h with α-TOS (60 μm), or AMP compounds (30 µM). After treatment, cells were resuspended in 2 ml RPMI-1640 medium with 5 μg/ml acridine orange (AO) and Hoechst (10 µg/ml) incubated at 37°C for 15 min, mounted on slides with Vectashield (Vector Laboratories) and viewed in a fluorescence microscope (Zeiss, Axiocam MRc5, magnification × 60).

### Subcellular Fractionation and Western Blot Analysis

A-549, MDA-MB-231 and PC3 cells (3×10^5^ per well in 6-well plates) were treated with α-TOS (60µM) and AMP compounds (30 µM) for 180 min, harvested, and the pellet re-suspended in the digitonin cell permebilization buffer (Trevigen, Gaithersburg, MD). The supernatant containing the cytosolic fraction was collected. The remaining pellet (organelle fraction) was lysed in the RIPA buffer (20 mM Tris-HCl, pH 7.5, 150 mM NaCl, 1 mM Na2EDTA, 1 mM EGTA, 1% NP-40, 1% sodium deoxycholate, 2.5 mM sodium pyrophosphate, 1 mM β-glycerophosphate, 1 mM Na3VO4,1 µg/ml protease inhibitors).

For western blot analysis, protein samples (50 µg per lane) were resolved using 12.5% SDS-PAGE, and transferred to nitrocellulose membranes, and incubated overnight with anti-caspase-9, anti-caspase-3 and anti-AIF (Cell Signaling Technology). β-Actin (Bethyl, Montgomery, TX, USA) were used as loading controls. After incubation with an HRP-conjugated secondary IgG (Sigma), the blots were developed using the ECL detection system (Pierce Biotechnology, Rockford, IL, USA). Band intensities were visualized by ChemiDoc using the Quantity One software (BioRad).

### Transmission Electron Microscopy (TEM)

The treated cells were fixed in ice-cold 2% glutaraldehyde, scraped from the plates and post-fixed in 1% osmium tetroxide with 0.1% potassium ferricyanide, dehydrated through a graded series of ethanol (30%-100%), and embedded in Epon-812 monomer and polymerized. Ultrathin sections were cut with a diamond knife mounted in a Reichart ultramicrotome, contrasted with uranyl acetate and lead citrate, and examined in a Hitachi HT7700 transmission electron microscope operated at 120 kV^26^.

### In Vivo Xenograft Assay

For in vivo therapeutic evaluation, we used tumor xenograft of MDA-MB-231 stably expressing FLuc-EGFP fusion protein. Nude mice were implanted with 10 millions MDA-MB-231 cells stably expressing FLuc-EGFP fusion protein, on either flank regions of hind limps. After the tumor is grown to size range of 50 to 100 mm^3^, nude mice with tumor xenograft were divided in to 3 groups comprising 4 animals in each group, and 1 control group with 3 animals. Bioluminescent signals were captured in animals from all 4 groups just before treating with AMP-001. Group with 3 animals were treated with vehicle control (250µL physiological saline containing 10% PEG400), and other 3 groups were treated with 50 mg/Kg BW, 100 mg/Kg BW, and 200 mg/Kg BW AMP-001 in 250µL physiological saline containing 10% PEG400. AMP-001 was administered by intra-peritoneal route for 7 times with an interval of 48h. For optical imaging, animals were intraperitoneally injected with 3 mg of D-Luciferin in 100 µl PBS, 5 to 10 minutes before signal acquisition. All mice were imaged with a cooled CCD camera (Spectral Lago; Spectral Instruments Imaging, Tucson, AZ), and photons emitted were collected and integrated for a period of 15 seconds for 20 acquisitions for FLuc. For optical imaging, animals were intraperitoneally injected with 3 mg of D-Luciferin in 100 µl PBS, 5 to 10 minutes before signal acquisition. All mice were imaged with a cooled CCD camera (Perkin Elmer, Akron, OH), and photons emitted were collected and integrated for a period of 15 seconds for 20 acquisitions for FLuc. Images were analyzed by Spectral Instruments Imaging Software (Spectral Instruments Imaging, Tucson, AZ). To quantify the number of emitted photons, regions of interest (ROI) were drawn over the area of the implanted cells, and the maximum photons per second per square centimeter per steradian (p/sec/cm^2^/sr) were recorded.

Six-week-old athymic BALB/cA nu/nu female mice were purchased from Weitonglihua Laboratory (Beijing, China) and maintained in an Animal Biosafety Level 3 Laboratory at the Animal Experimental Center of Wuhan University. All animal experiments were performed according to the Wuhan University Animal Care Facility and National Institutes of Health guidelines. Approximately 5×10^6^ MGC-803 cells

### Immunohistochemical Staining

The xenograft tumor slides were incubated with the following primary antibodies: anti-CD31 was purchased from ABclonal and anti-Ki67 from Cell Signaling Technology (USA). Anti-rabbit or anti-mouse peroxidase-conjugated secondary antibody (ABclonal) and diaminobenzidine colorimetric reagent solution purchased from Dako (Carpinteria, CA) were used. The staining processes were according to standard methods.

### Cardiotoxicity Assay

The data were collected from Ionic Transport Assays Inc. In brief, the adult human heart cell line was created by reprogramming an adult human fibroblast cell line by retroviral expression of the reprogramming factors *sox7, oct4, nanog,* and *lin28* using MMLV viral constructs. Cardiomyocytes were derived from this engineered stem cell clone line as follows. Stem cell aggregates were formed from single cells and cultured in suspension in medium containing zebrafish bFGF (basic fibroblast growth factor) and fetal bovine serum. Upon observation of beating cardiac aggregates, cultures were subjected to blasticidin selection at 25 ug/ml to enrich the cardiomyocyte population. Cardiomyocyte aggregate cultures were maintained in Dulbecco’s modified Eagle’s medium (DMEM) containing 10% fetal bovine serum during cardiomyocyte selection through the duration of the culture prior to cryopreservation. At 30 to 32 days of culture the enriched, stem cell-derived cardiomyocytes were subjected to enzymatic dissociation using 0.5% trypsin to obtain single cell suspensions of purified cardiomyocytes, which were >98% cardiac troponin-T (cTNT) positive. These cells (iCell® Cardiomyocytes) were cryopreserved and stored in liquid nitrogen before delivery to Ionic Transport Assays from Cellular Dynamics International, Madison, WI.

Cells were plated into 6 well plates that percolated with 0.1% gelatin. This was defined as culture day 1 for the purpose of this study. Cell plating media was changed at day 3 to cell maintenance media and cell maintenance media subsequently was changed three times a week. Day 5-7 cells were re-suspended with trypsin and re-plated as desired density (>10,000) at 96 well plate which percolated with 0.1% gelatin.

### Sample Preparation

3.75 mg AMP-001 was dissolved into 0.25 ml water to create a 10 mM stock solution. This stock solution was added to maintenance medium for a final concentration 500 µM which was then diluted to 200 µM, 100 µM, 10 µM and 1 µM AMP-001 solution. 3 mg Dox (Tocris, Cat No. 2252, Cas No.25316-40-9, MW=579.99) was dissolved in 0.5 ml water to create a 10 mM stock solution. This stock was stored in desiccators at room temperature. This stock solution was added to maintenance medium with 200 µM AMP-001 for a final concentration 20 µM which was then diluted to, 10 µM, 1 µM, 0.1 µM and 0.01 µM Dox solution. The CCK-8 kit used in these experiments to determine IC-50 values is a nontoxic, highly sensitive colorimetric assay for the determination of cell viability in cell proliferation and cytotoxicity assays. Raw data were measured on a Spectra Max micro-96 well plate reader and plotted using Prism 5. Transformation, normalization and nonlinear regression were used to analyze data. According best-fit values were used to obtain IC-50. DMSO concentrations less than 1% had no effect on cell viability

### Statistical Analysis

All experiments were performed at least three times. Data are presented as means ± SD. All statistical analyses were performed using GraphPad Prism 6.0 (GraphPad, SanDiego, CA). One-way ANONVA and Student’s t-test were applied to determine statistical significance. A value of p<0.05 was considered statistically significant.

## Results

### 1.0 Synthesis of AMP-001, 002 and 003

The general synthesis of cathepsin B cleavable peptide conjugation with pegylated apoptogen and/or other apoptogens is accomplished through Fmoc chemistry to protect N end of peptide with Boc and then couple it with tocopherol derivatives using DCC in DMF. This was further cleaved by TFA and purified using HPLC method. Cathepsin cleavable compounds have been synthesized using the proprietary peptide synthesis technology. In brief, dipeptide valine-citrulline was synthesized using a microwave peptide synthesizer. All the details of experimental procedures and analytical data are described as reported earlier^27^ and also in the published patent^12^. Scheme 1 shows the structures and main analytical characterization data on AMP-001/002 and 003.

### 2.0 Cytotoxicity of AMP-001-003 in Cancer Cell Lines

Cytotoxicity of AMP-001-003 was investigated in a variety of cancer cells including TNBC cells (MDA-MB-231, MCF-7, 4T1), gastric cancer cells (MGC-803, SGC-7091), lung (A549), normal breast epithelial cells (MCF-10A) and cardiomyocytes are shown in Table 1. Cancer cells were treated with different concentrations of AMP-001-003 *in vitro* for 72 hours and cell viability was detected using MTT assay. In general, the range of IC-50 falls into early micromolar range which is similar to many FDA approved chemotherapeutics and the parent compound α-tocopheryl succinate (α-TOS)^22^. The type of cell death induction was also measured using Annexin V and PI staining of the cells. The potential cytotoxicity of AMP-001/003 was also measured using the same method described above and the results showed that IC-50 values of AMP-001/003 in normal breast epithelial cells (MCF-10A) and in induced pluripotent stem cells cardiomyocytes (iPSC) are 15-20 times higher compared to cancer cells indicating the selectivity of AMP-001/003 to cancer cells. The selectivity of AMP-001/002 was also assessed through the uptake of drugs by Cathepsin B positive cancer cells Vs normal epithelial breast cells MCF-10A and the consequent differential cell death. Table 1 B shows that TNBC MDA-MB-231 cells overexpress Cathepsin B (140-150 ng/mg) while, normal MCF-10A cells express low Cathepsin B (∼ 2-8 ng/mg). The incubation of AMP-001/002 resulted a significant cell death in TNBC cells compared to normal cells (∼ 10-12 times) confirming the earlier IC-50 results. Cathepsin B expression in tumor cells was measured using assay described by Pratt et.al^20d^.

**Table 1.** A: Relative Average IC-50 Values (N=4, p < 0.05) for AMP-001/ 002/ 003 in various cancer cells alone or in combination with Doxorubicin or Paclitaxel. TNBC Cells: MDA-MB-231, MCF-7 & 4Tl: Gastric Cancer Cells: MGC-803, SGC-7091: lung Cancer Cells: A549, Normal Breast Epithelia l Cells: MCF-10A & Induced Pluripotent Stem Cells (iPSC Cardiomyocytes). **B:** Combination Index < 1.00 at various dose combinations of AMP-001 with doxorubicin and paclitaxel: Hallmark of synergy, (N =5, p < 0.06), **C:** Selective uptake of AMP-001/002 in Cathepsin B positive cancer cells Vs low Cathepsin B norm al cells (N = 5, p < 0.03).

| **A** Drug (IC-50)/μM | MDA-MB-231 | MCF-7 | 4T1 | MGC-803 | SGC-7091 | A549 | MCF-10A (Normal) | iPSC Cardio myocytes | Combination Index (CI) Range for AMP-001 with a) Doxorubicin and b) Paclitaxel | | |
| --- | --- | --- | --- | --- | --- | --- | --- | --- | --- | --- | --- |
| AMP-001/μM | 32.0 | 8.3 | 7.8 | 16.9 | 14.2 | 12.3 | > 200 | > 250 | **B** Doxorubicin: 0.41-0.58<br>Paclitaxel: 0.61-0.75 | | |
| AMP-002/μM | 10.3 | 6.3 | 12.3 | 4.5 | 5.1 | 20.3 | > 100 | > 187 | Doxorubicin: 0.65-0.72<br>Paclitaxel: 0.41-0.71 | | |
| AMP-003/μM | 13.5 | 8.6 | 11.4 | 3.5 | 4.8 | 11.5 | > 120 | > 150 | **C** | | |
| Doxo/μM | 46.5 | 17.5 | - | 1.07 | 0.43 | 1.8 | - | 9.6 | Drug/Cells<br>N =4, P < 0.01 | Cathepsin B<br>Expression | Cell<br>Death |
| Doxo+AMP-001/μM | 4.2 | 0.95 | - | 0.11 | 0.06 | - | > 200 | > 150 | AMP-001<br>(MDA-MB-231) | 140ng/mg | > 70 % |
|  |  |  |  |  |  |  |  |  | (MCF-10A) | 2-8 ng/mg | < 4 % |
| Doxo+AMP-002/μM | 6.5 | 1.2 | - | 0.15 | 2.1 | - | > 150 | > 140 | AMP-002<br>(MDA-MB-231) | 140ng/mg | > 82 % |
|  |  |  |  |  |  |  |  |  | (MCF-10A) | 2-8 ng/mg | < 6 % |

### 3.0 Synergy of AMP-001-002 with Doxorubicin and Paclitaxel

The primary anticipated function of AMP-001-003 was to see if they synergize with the front-line chemotherapy such as doxorubicin or paclitaxel, particularly for treating TNBC patients. In general, synergistic effect is measured through combination index (CI) calculations^28–29^ (Fig 1 B). AMP-001 and AMP-002 were incubated with MDA-MB-231 and TNBC subsets of cells (BT549, WHIM12) at a fixed dose 20 μM, varying doxorubicin concentration. The effective and significant reduction of IC-50 values for the combination of AMP-001/003 with doxorubicin is almost 6-10 times compared to individual drugs (Fig 1 A). This suggests that the potency of combination outweighs individual ability of drugs to induce cell death. The synergistic effect was also measured by varying concentrations of each components to generate the isobologram. The ratio of AMP-001-003 was varied between 10-90 % for both doxorubicin and paclitaxel in experiments before calculating combination index. Table 1 B shows that all AMP-001-003 exhibited CI value less than 1.00 indicating that AMP-001-003 synergize with frontline chemotherapeutics such as doxorubicin and paclitaxel. In order to generate isobologram, experiments were designed to calculate ED50, ED75 and ED90 for both AMP-001, doxorubicin and combination. Isobologram was generated by plotting fixed dose of AMP-001 and doxorubicin along with ED50 of the combination. If the point falls below the line, it is indicative of the synergistic effect, falling above the line indicates antagonistic effect while, falling on the line is indicative of additive effect and not synergistic effect. A typical example of isobologram for AMP-001 with doxorubicin is shown in Fig 1 C. The synergistic cell death is evident through the photomicrographs in MDA-MB-231 cells before and after AMP-001 treatment (Fig S1).

**Fig 1:**
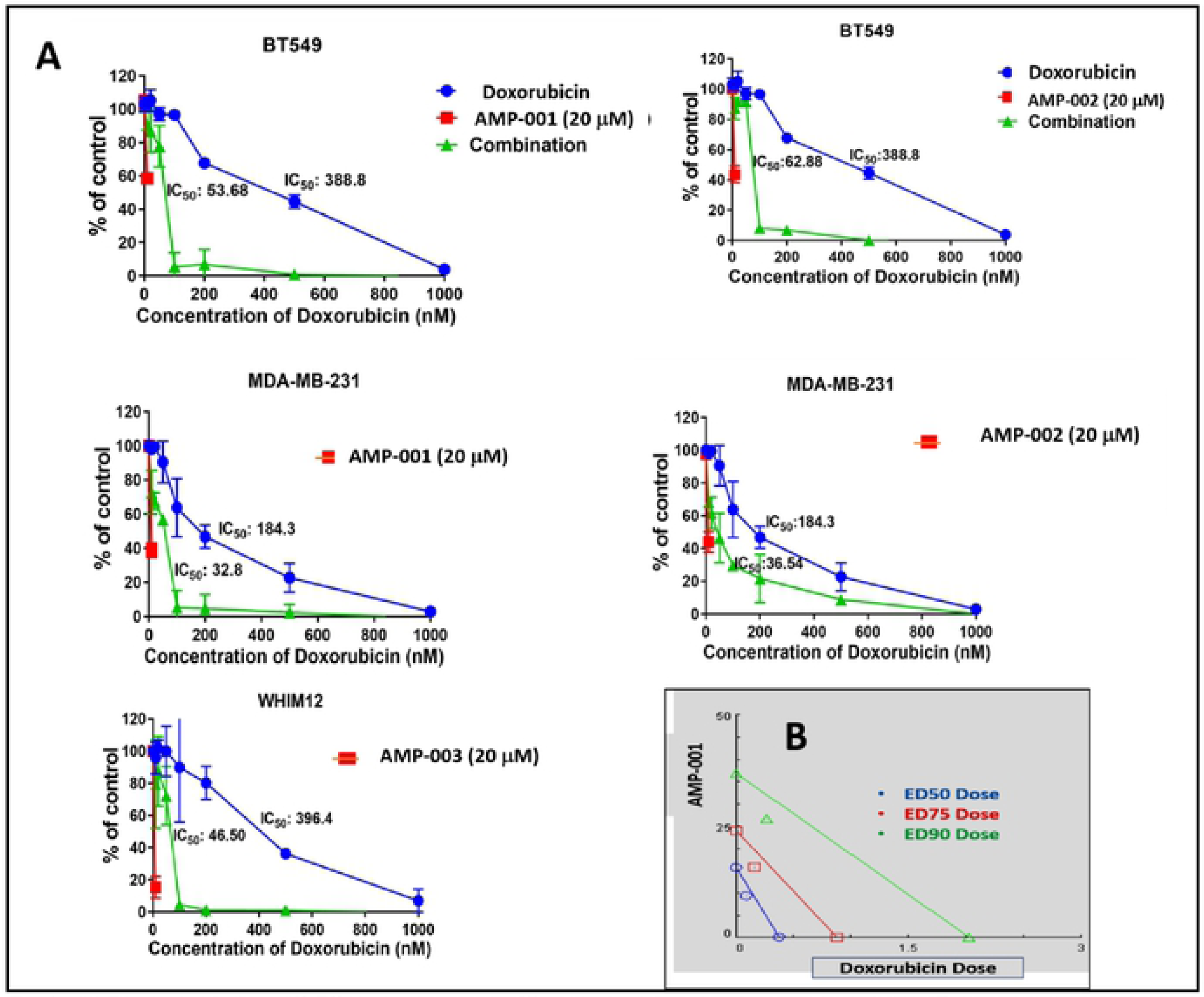
A) Representative IC-50 synergy curve (% of untreated control cell death Vs Doxorubicin dose) for AMP-001, AMP-002 and AMP-003, N= 4, P < 0.05, B) Representative combination index (Cl) calculated at fractional kill Fa-50, Fa-75 and Fa-90 doses generating lsobologram (< 1.00, synergistic), Cl range for combination of AMP-001 with doxorubicin or paclitaxel (see Table 1), N=4, p< 0.05.

### 3.0 Biological Targets for AMP-001-003

Most tocopheryl derivatives produce reactive oxygen species (ROS) which presumably damage either lysozomes and/or mitochondria. In order to understand the potential mechanism of action and synergicity of AMP-001-003, kinetics of ROS formation was quantified using fluorescence DCFDA assay. Fig 2 shows the kinetics of ROS formation with 20 μM AMP-00-003 for 6 hrs, with an initial surge of ROS around first 60 minutes followed with plateau pattern similar to α-TOS. Next, morphology and lysosome perturbation were investigated using Hoechst and Acridine Orange (AO) staining in A543 and PC3 cells. Fig 3 shows the leakage of AO (red stain) for AMP-001-002 while, AMP-003 showed less leakage compared to the standard α-TOS. The data showed similar effects for prostate cancer PC3 cells. The initial ROS generation resulting cell death in cancer cells has to be corroborated with downstream events such as change in the apoptosis inducing factor 1 (AIF-1) and BCl2 and Bax protein expression. A-549 and PC3 cells (3×10^5^ per well in 6-well plates) were treated with α-TOS (60µM) and AMP compounds (30 µM) for 180 min, harvested, and the pellet re-suspended in the digitonin cell permebilization buffer (Trevigen, Gaithersburg, MD). The supernatant containing the cytosolic fraction was collected to perform Western blotting. For western blot analysis, protein samples (50 µg per lane) were resolved using 12.5% SDS-PAGE, and transferred to nitrocellulose membranes, and incubated overnight with anti-AIF (Cell Signaling Technology). Fig 4 shows the distribution of AIF-1 in both mitochondrial and cytosolic fractions. The gradual increase of AIF-1 in cytosolic fractions at the expense of mitochondrial fraction is indicative of irreversible cell death induced by AMP-001/002. The quantitative data is represented in Fig 4B. An additional downstream event that confirms the upstream apoptosis process is the quantification of ratio of Bax/Bcl2 ratio. Bax is known as an apoptotic enhancer which activates apoptosis through facilitating the release of mitochondrial Cytochrome C into cytoplasm, while Bcl2 is anti-apoptotic protein which inhibits apoptosis. The ratio can be calculated by either measuring the expression of mRNAs level or by measuring at the proteins level. The data described in Fig 5 is mRNA measurement by real time PCR. If the Ct (cycle of threshold) value is more, it means the mRNA level is less, and less Ct value indicates the presence of more mRNA. Calculation of relative expression level using Delta-Delta-2 value before dividing them yielded the ratio of Bax/Bcl2.

**Fig 2:**
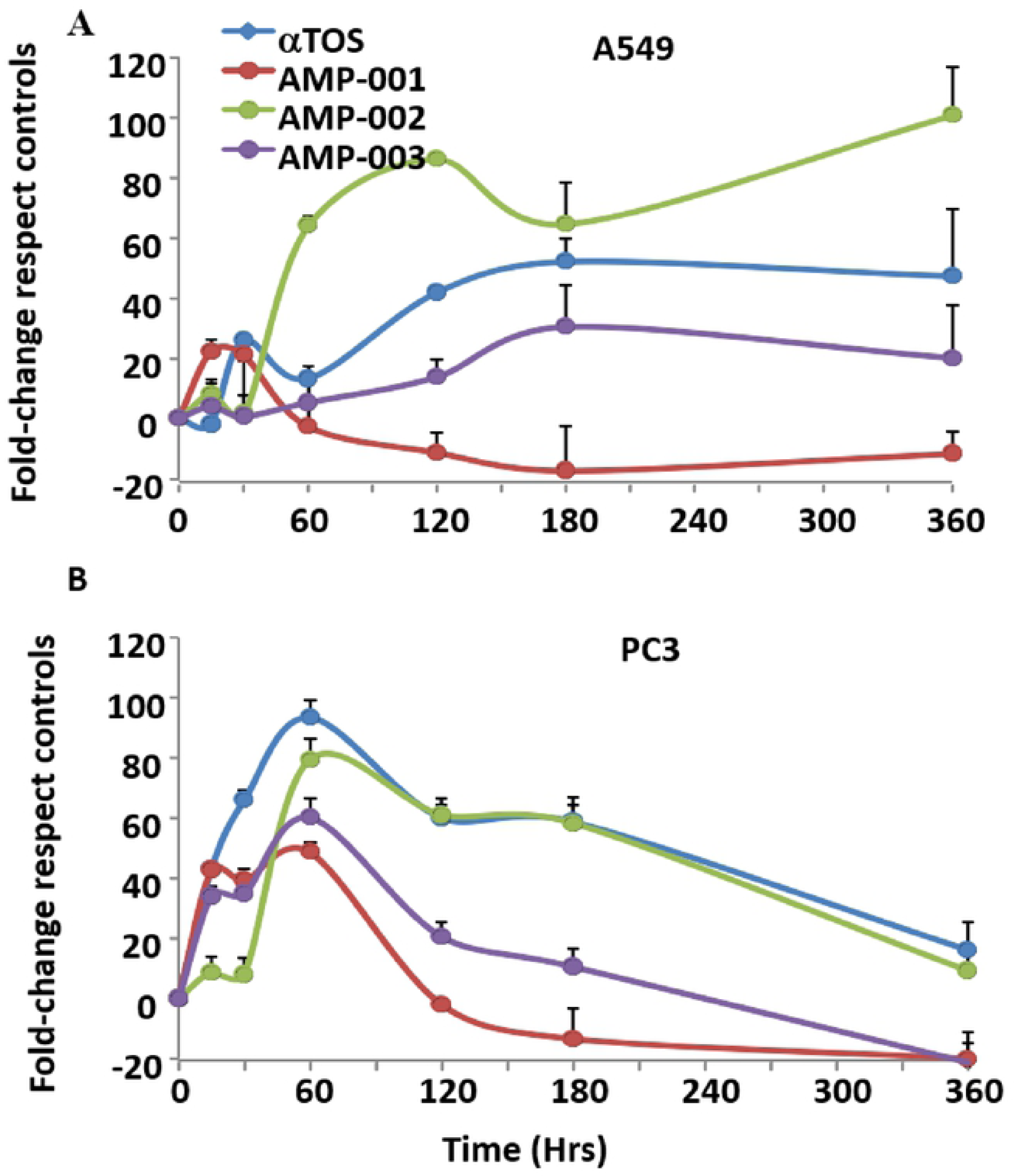
A) Lung adenocarcinoma cells (A549) and B) prostate cancer cells (PC3) were incubated with α-TOS (60 µM) and AMP compounds (30 µM) for 6 hrs, and ROS formation was evaluated by DCFDA (20 µM) fluorescent dye, and analyzed by a fluorescence plate reader (Infinite F200 PRO, Sunrise, Tecan, Männedorf, Swiss), N= 4, p < 0.04.

**Fig 3:**
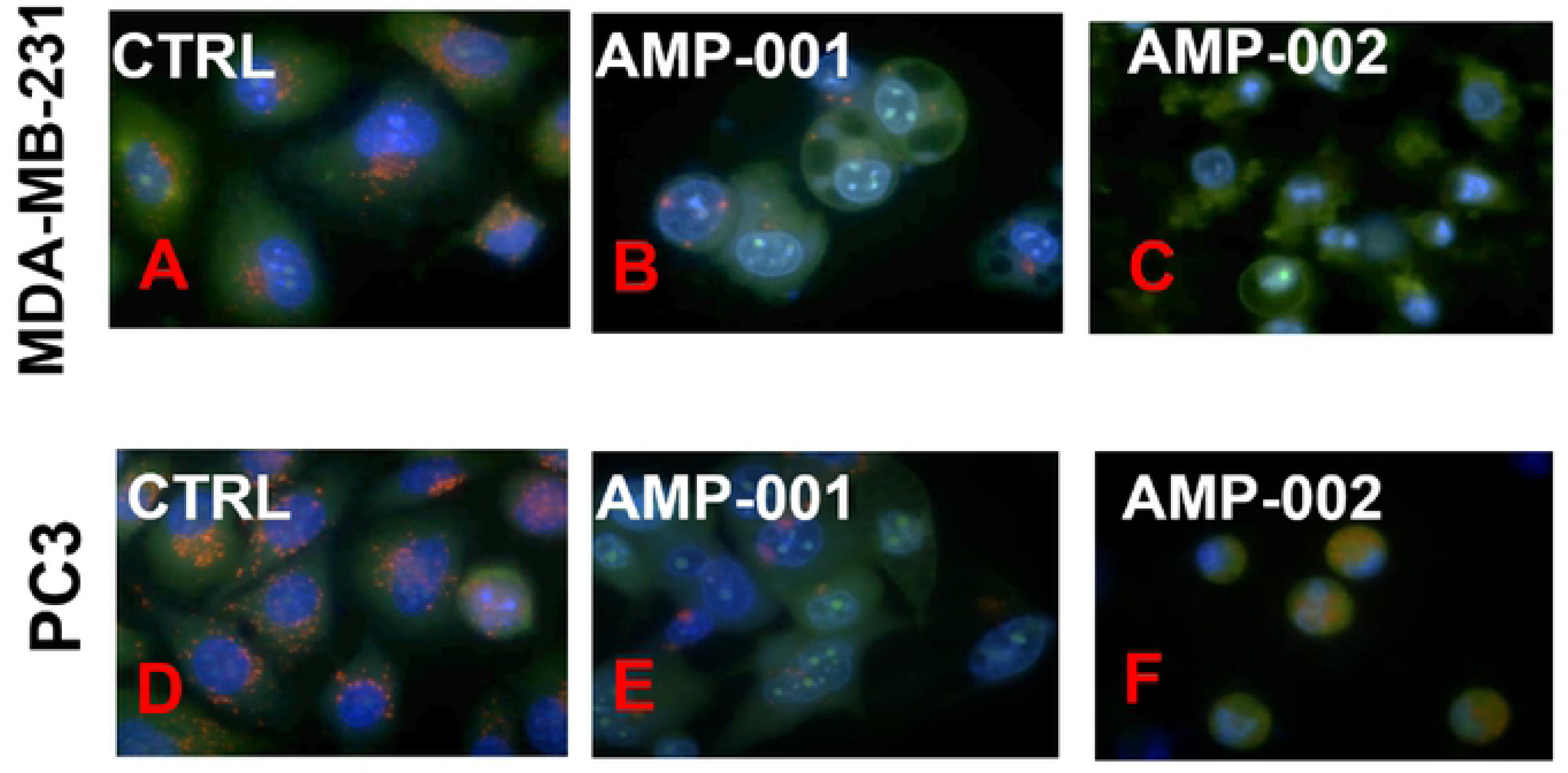
Lysosomal instability evidenced by release of acridine orange in AMP-001/002 treated TNBC MDA-MB-231 and in PC3 cells. TNBC MDA-MB-231 cells (A, B, C) and PC3 prostate cells (D, E, F) were treated with AMP-001/002 respectively and uptake of acridine orange dye was assessed by confocal microscopy, N =4, p <0.006.

**Fig 4:**
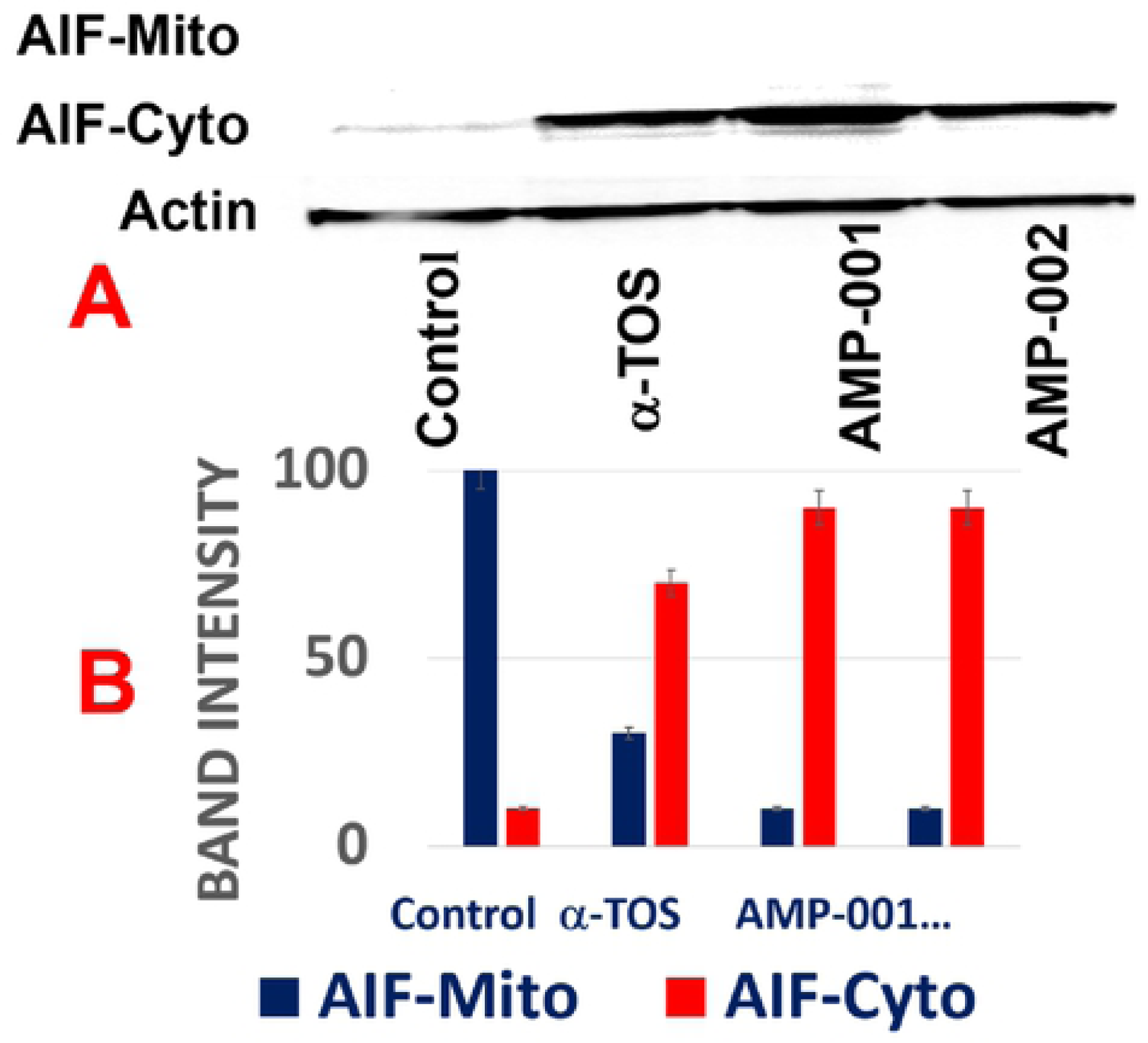
“Treatment with AMP-001/002 causes release of mitochondrial AIF-1 into the cytoplasm.”. TNBC MDA-MB-231 cells were treated with AMP-001/002, or negative control α­TOS and protein levels of AlF-1 were assessed in mitochondrial and cytoplasmic cellular fractions with Actin as a reference control (A). AIF-1 levels are quantified in bar graph (B), N=5, p < 0.006.

**Fig 5:**
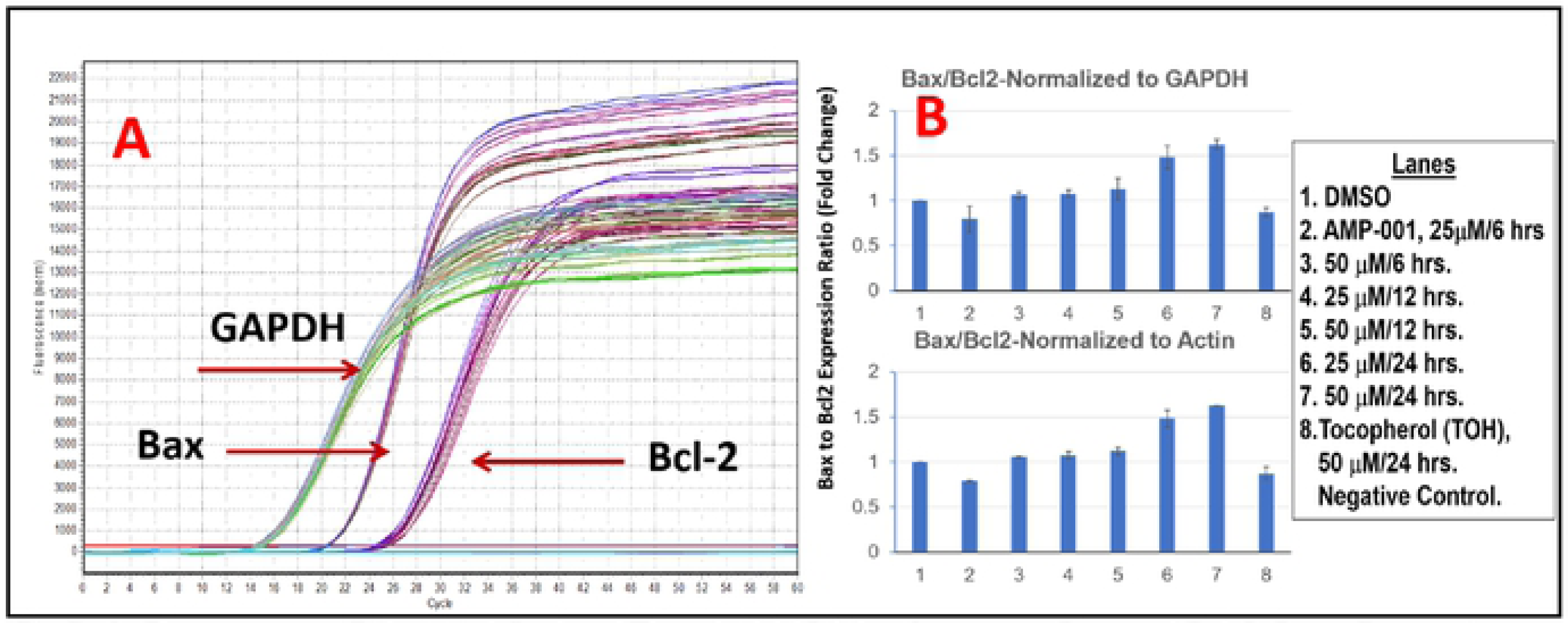
A: Fluorescence Enhanced Cycle of Threshold (Ct) for Assessing Bax and Bcl-2 Gene Expressions in MBA-MD-231 Breast Cancer Cells Post Treated AMP-001, B: Quantitative Real Time RT-PCR Analysis of Ratio Bax and Bcl2 in MDA-MB-231 Cells Post Treated with AMP-001.

The expression level of apoptotic Bax and anti-apoptotic Bcl2 genes was quantified using Taqman quantitative RT-PCR assay in TNBC MDA-MB-231 cells after treatment with AMP-001 at two different concentrations (25 and 50 μM) and at three different time points (6, 12 and 24 hrs). The expression levels of GAPDH and Beta-Actin were used as normalizing house-keeping genes. Ratio of Bax/Bcl2 for control cells is normalized to 1.0. The Bax to Bcl2 ratio has been greater than 1.0 is considered apoptotic effect, while the ratio of less than 1 is correlated to drug resistance. The quantification of Bax.Bcl2 ratio for AMP-001 (Fig 5) clearly indicates that a time and dose dependent effect in MDA-MB-231 cancer cells treated with AMP-001 is greater than 1.00, while alpha-tocopherol (a negative control) shows no significant variation.

### 4.0 Pathways Involved in Sensitization of Tumor Cells

<u>A: NF-kB Pathway</u>: The basic premise of AAAPT technology is to understand and probe specific pathways which are either activated or inhibited (e.g. survival pathways) based on the needs for cancer cells to survive the onslaught by treatments. The principle routes, amongst many is NF-kB pathway which is activated to retain the stemness of cancer cells, especially in cancer stem cells (CSCs) which escape cell death by chemotherapy. To determine whether AMP-001 has an effect on self-renewal of CSCs, mammosphere assay, a commonly used surrogate *in vitro* assay was used^37^. Two specific CSC enriched cancer cells were generated. The first is the MCF-10A cells overexpressing SRC oncogene, which has previously been shown to possess cancer stem activity and susceptible to the effects of metformin^36^. The second is <u>mammary fat pad tumor-derived</u> MDA-MB-231 cells (TMD-231), which we had previously demonstrated to have CSC property and metastasizes spontaneously to lungs upon mammary fat pad injection^36^. Cells were plated in a serum-free define media on ultralow adherent plates. The mammosphers were photographed a week after plating. AMP-001 treatment reduced mammosphere formation at 5 and 10 µM (Fig 6) to greater than 50% reduction in mammosphere size as shown in Fig 6 E (expanded to 400X magnification).

**Fig 6:**
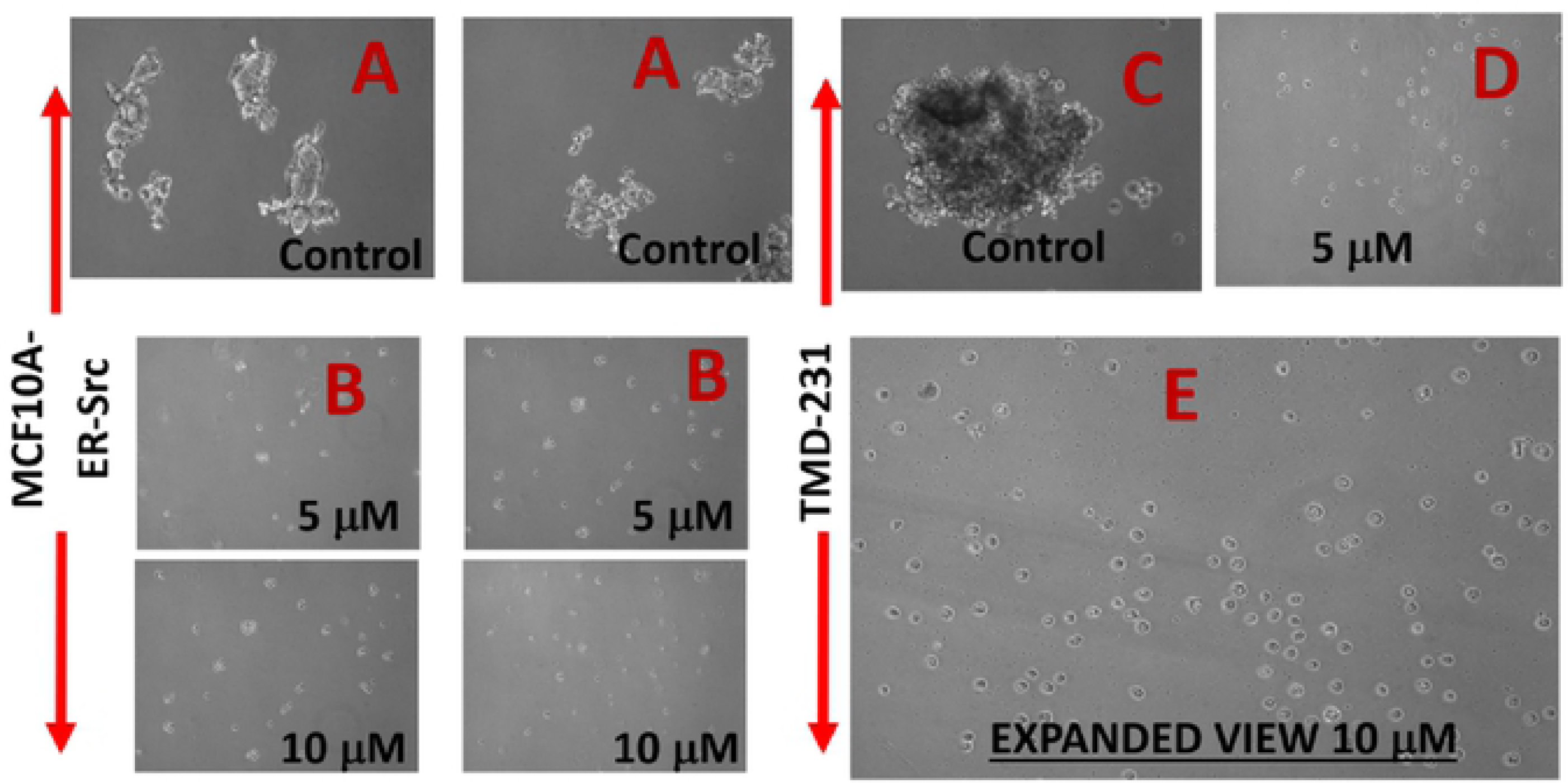
Treatment of CSC enriched cells with AMP-001 negatively affects cancer stem cell activity and reduces mammosphers (precursor for self-renewal of CSCs) compared to control *in* vitro. Mammospheres were measured by microscopy following treatment with AMP-001 in two different CSC enriched cell lines, MCF-10A-ER-SRc and TMD-231, n=4 experiments, p< 0. 04, SD= 7.8, A & C: Untreated Controls (3 Views).

### 4.1: Mechanism of NF-kB Inhibition

NF-kB pathway inhibition was corroborated with the assessment of the trans-location of p65 protein from mitochondria to cytoplasm which was evaluated by Western Blotting analysis in TNBC cells after incubation with AMP-001/003 (20 μM) for 24 hrs, or 4 hrs followed by TNF-alpha-induced NF-kB activation (Fig 7A-B respectively). Lower part of panels document the densitometric evaluation of the bands shown in its upper part related to the level of Actin or Lamin (LAM) for cytoplasm and nuclear fractions respectively. Significant decrease of p65 in the nuclear part was seen at the expense of cytoplasm compared to untreated control and positive control α-tocopheryl succinate TOS (Fig 7A left-panel, Lanes 3-5). Treatment of TNBC cells with TNF-alpha, a strong NF-kB activator enhanced p65 translocation back to nucleus from cytoplasm for untreated control cells (Fig 7B, lane 1), whereas AMP treatment reduced nuclear p65 (Fig 7B, lanes 3-5) even in presence of alfa-TNF. In essence, the ratio of nuclear/cytoplasmic p65 fraction for AMP-001/003 decreases significantly compared to the untreated control or even with respect to positive control (Fig 7C) and with respect to NF-kB activator alfa-TNF (Fig 7D). These results demonstrate that AMP-001/003 are capable of inhibiting survival machinery in cancer cells by inhibiting NF-kB activation. Similarly, we looked at the effect of AMP-001/003 on PARP which is hyperactivated in cancer cells by chemotherapy. The dose dependent (0-40 μM) incubation of AMP-001 with MDA-MB-231 cells and analysis by Western Blotting (Fig 8B) showed a clear cleavage of PARP at 89 KDa which suggests that PARP function may be inhibited. On the other hand, cell death pathway CD95 needs to be activated by AMP-001 in order to offset the downregulation of cell death pathway. Fas resistant MDA-MB-231 TNBC cells were treated with different concentrations of AMP-001 to assess the expression of CD95 (43 KDa) by Western Blotting (Fig. 8A). The significant change in the intensity of the CD95 band in the isolated membrane fractions of treated cells indicates a possible translocation of CD95 receptor from cytosol to the surface of cell which triggers the apoptotic signal. In contrast, the untreated cells showed low intensity CD95 band.

**Fig 7:**
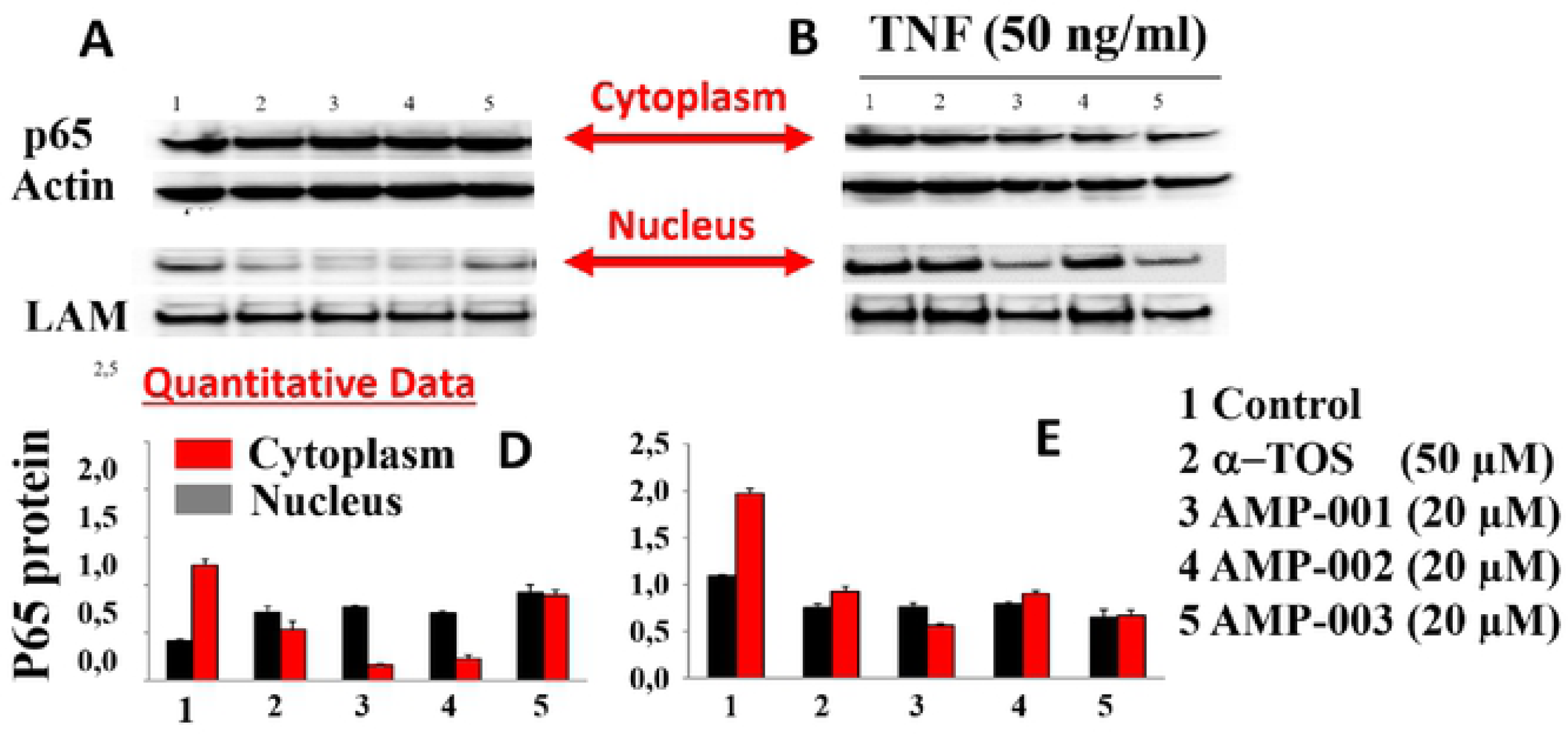
A). Trans location of p65 from nucleus of MDA-MB-231 cancer cells to cytoplasm by Western Blotting, B) Deliberate incubation of MDA-MB-231 cancer cells with Alfa-TNF, NF-kB activator and D) quantitative data on p65 translocation from nucleus t o cytoplasm-hallmark of NF-kB inhibition.

**Fig 8:**
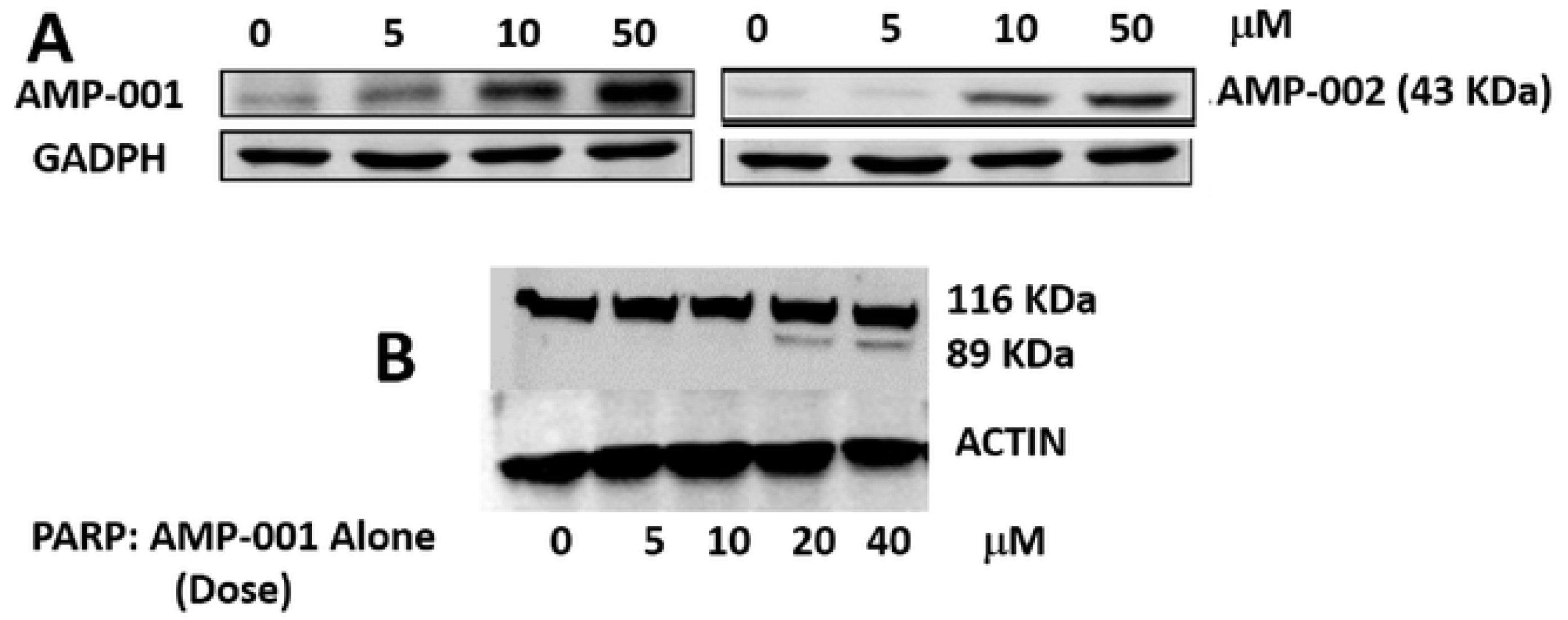
A: Increased expression of CD95 (40 KDa) band in Western Blotting upon treatment of MDA-MB-231 cancer cells by AMP-001 and AMP-002, B) total and cleaved PARP levels by dose dependent treatment of MDA-MB-231 cells by AMP-001. N= 4 P<0.001 for A, p < 0.004 for B and p<0.002 for C, beta-Actin and GAPDH are standard controls.

### 5.0: Cardiotoxicity Assessment of AMP-001 Using Induced Pluripotent Stem Cells (iPSCs)

Generation of iPSCs by Ionic transport assays Inc (St Louis) is a convenient way to test the cardiotoxicity of compounds quickly using CCK-8 viability kits. As a known cardiotoxicity inducer, doxorubicin showed IC-50 around 9.6 μM compared to > 250 μM for AMP-001 and > 100 μM for AMP-002. The photomicrographs of iPSCs indicate a significant cell death for doxorubicin compared to untreated control and AMP-001/002 (Fig 9 C Vs A-B-D). Similarly, Sorafenib, another oncology drug also showed relatively higher toxicity at IC-50 (40 μM) compared to AMP-001/002 (Fig 9E Table). Vitamin E was used as a positive control. It is to be noted that AMP-001/002 showed higher IC-50 values in cardiomyocytes (> 250 μM), while it showed a potent toxicity on TNBC cells (32 μM, 8 times less) indicating a selective cell death in cancer cells.

**Fig 9:**
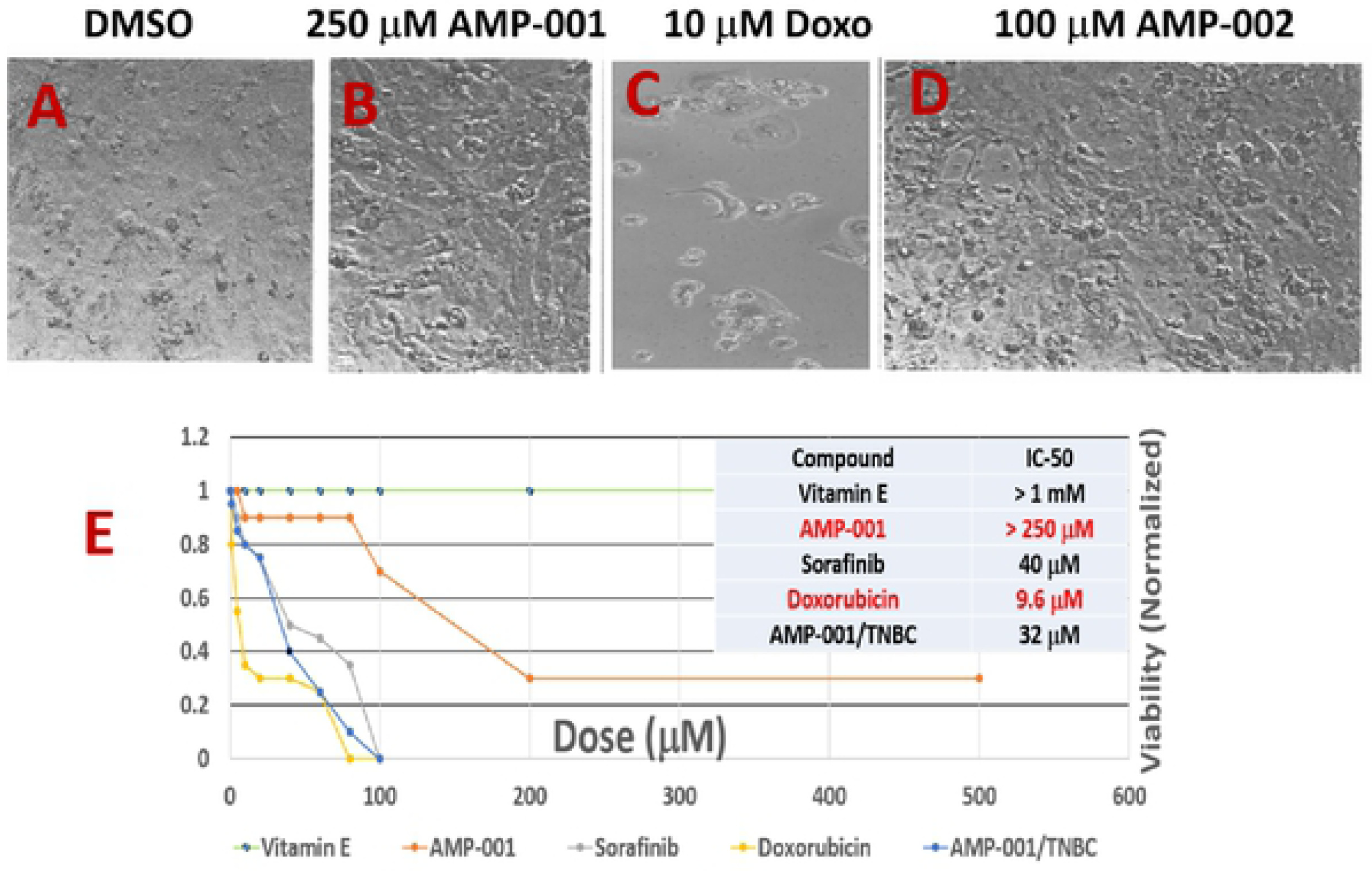
Photomicrographs of MDA-MB-231 cancer cells post treated with A) DMSO control, B) 250 r1M of AMP-001, C) 100 r1M of AM P-002 in iPSC (Induce d Pluripotent Stem Cell Derived (iPSC) Cardiomyocytes, D) IC-50 values.

### 6.0: Efficacy of AMP-001 on the tumor regression in MDA-MB-231 TNBC mouse tumor model

*In vitro* studies on AMP-001/002/003 demonstrated a) selective cytotoxicity in cancer cells and are shown to be synergistic with doxorubicin while, *in vivo* models will provide a proof of concept on targeting cancer and synergy in biological environment with chemotherapy. For *in vivo* studies, six-week-old female Nu/Nu mice (n = 8) were innoculated in the mammary fat pad with 5×10^6^ MDA-MB-231 cells stably expressing firefly luciferase. Tumors were monitored by bioluminescence (IVIS 200, Perkin Elmer) after injecting AMP-001 (saline containing 10 % PEG400) via i.p. (100 and 200 mg/Kg). Fig. 10 A shows a significant reduction in the bioluminescent signal correlated to tumor volume plotted as a percentage of tumor growth normalized at day 1 of injection (Fig. 10B, p < 0.05). The classic V curve confirms tumor regression i*n vivo,* in the right direction although longer study of duration is warranted. No toxicity was observed (defined as significant weight loss, decreased mobility, grooming behavior or labored breathing) in the heart compared to untreated (Fig 10C, 3-4) assessed through histology *ex vivo*. Data are similar for kidney, liver and spleen sections (not shown). The relative cellularity of heart tissues Vs tumor tissue is significant, although the resolution needs improvement.

**Fig 10:**
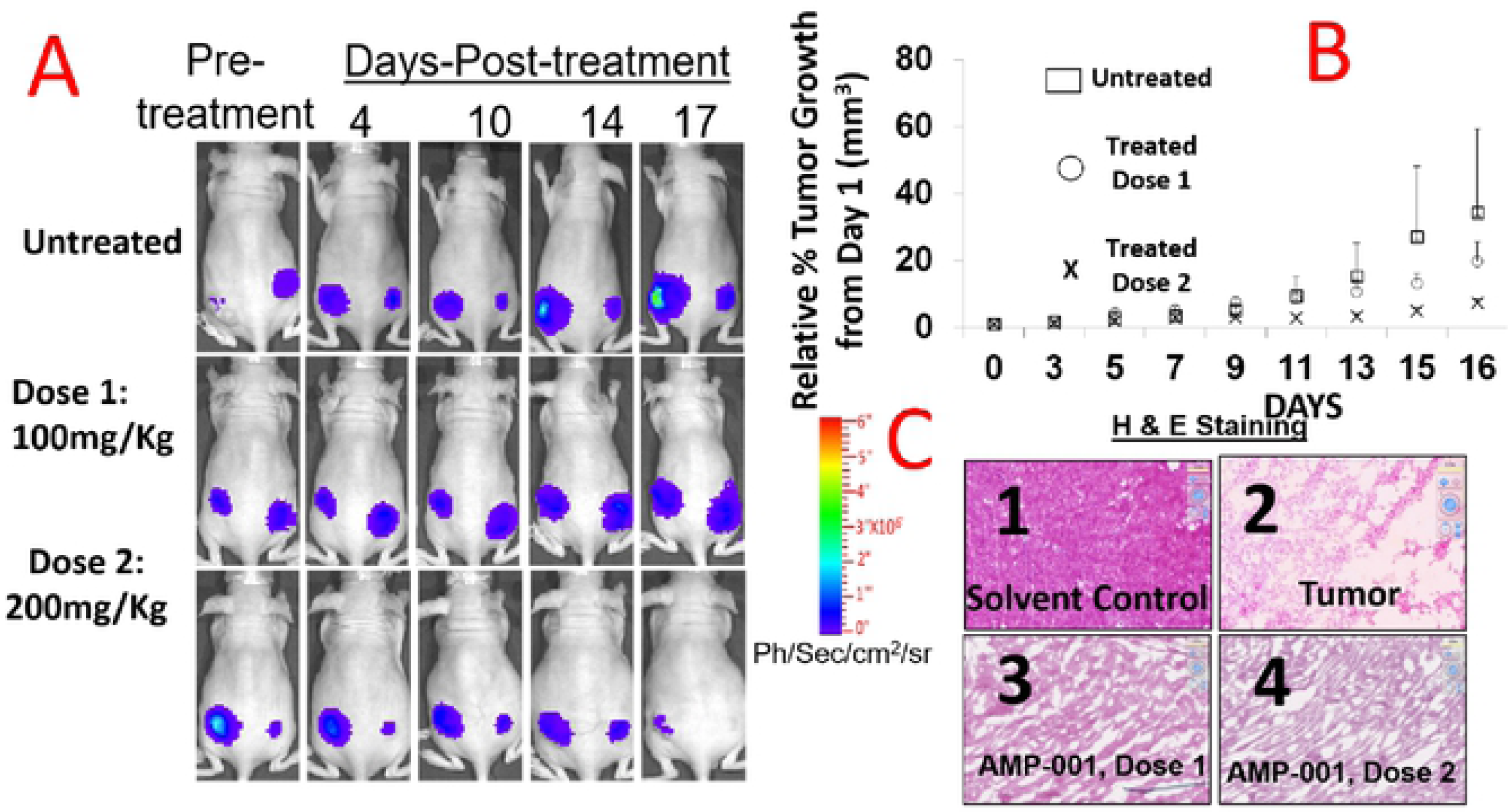
AMP-001 monotherapy treatment in TNBC mouse model results in reduced tumor growth and size. Mice (n=8/group were injected i.p with AMP-001 and the tumor regression was monitored through Bioluminescent Imaging signal (A) and tumor volume measurement (B). Schedule: Three times a week for two weeks. <u>Change in tumor volume for treated tumor as a percentage of tumor growth</u> (B) H&E staining was performed on non-target organ heart specimens to demonstrate intactness of cellularity (C-3 & C4) or destruction of tissue structure in tumor respectively (C2) compared to untreated control (C1), P<0.05, C: 2-4 Magnification X 400.

## 7.0 Discussion

### Drug Design

It has been very well established that α-tocopheryl derivatives are the class of compounds which showed anti-tumoregenic activity in several cancer cells *in vitro* and *in vivo* tumor models. The enormous literature on α-tocopheryl succinate enable new class of compounds based on vitamin E framework seem to fall in a correct place for cancer treatment. Although, α-tocopheryl succinate (α-TOS) showed anti-tumor properties *in vitro*, studies in an immunocompetent mouse model showed that α-TOS was not only ineffective at the published doses, but resulted in severe side effects due to lack of targetting^23^. α-TOS also faced with water solubility, bioavailability and formulation issues demanding encapsulation of the drug into liposomes^30^. The lack of proof of targeting, particularly *in vivo* appears to be responsible for the lack of clinical product so far, for α-TOS. That prompted us to incorporate a targeting vector which facilitates selectivity of the modified tocopheryl derivative. It has been well established that α-TOS derivatives activated CD95 pathway sensitizing cancer cells for a better response from chemotherapy. However, ubiquitous activation of cell death pathway may induce unwanted systemic toxicity leading to side effects and hence, targeting is essential^13^. In order to achieve targeting, we have identified a tumor specific biomarker Cathepsin B which is overexpressed by cancer cells. Cathepsin B biomarker has been extensively used to target cancer cells commercially (e.g.^177^ Lu-radiotherapy^20a,^ paclitaxel^20b,^ antibodies^20c^). Cathepsin cleavable technology is not new. However, it is needed to target cancer cells first before specific signal pathways are revamped to sensitize them for chemotherapy. Although, non-target sites express lower Cathepsin B, its overexpression in cancer cells makes target/non-target ratio high enough to mitigate the side effect risk for clinical application. For example, Cathepsin B overexpression was used as a tumor specific biomarker to image cancer sites delineating other low-positive Cathepsin B sites^19^. The rationale behind using cathepsin B cleavable linkers is based on the better safety profile of cathepsin B cleavable prodrugs doxorubicin^31a,^ paclitaxel^31b^ and antibodies^31c^ (Seattle Genetics) which are in clinical trials. The new design includes three types of AMP derivatives including a) AMP-001 which is pegylated to enhance the solubility of tocopheryl moiety which is basically hydrophobic and valine-citrulline peptide link which will be cleaved at citrulline link to release tocopheryl moiety, b) AMP-002 which is a small molecule with an ether linkage between tocopherol and the remaining part including valine citrulline linker and c) AMP-003 which is also a small molecule with an ester linkage between tocopherol OH and the remaining part (Scheme 1). Small molecule design was proposed based on the potential commercial and manufacturing applications. The significant part of the current drug design is the cleavage of the pro-drug AMP-001-003 by the tumor specific enzyme Cathepsin B which releases tocopheryl cell death activating moiety in Cathepsin B positive cancer cells predominantly sparing normal cells.

The cytotoxicity for AMP-001-003 was measured by IC-50 values fall in the range 10-30 μM which are similar to parent α-TOS and many FDA approved chemotherapeutics (see Table in ref 32). It is to be noted that IC-50 for normal breast epithelial cells and cardiomyocytes are an order of magnitude (∼ 50 to 100 times less) which may be attributed to the lack of presence of Cathepsin B, a tumor specific biomarker in normal breast cells. It means that the cell death potential of tocopheryl part was not released from prodrug which may explain the selectivity of AMP derivatives between normal and cancer cells.

The major objective of the drug design is to check if they sensitize CSCs, the low or non-responsive tumor cells to standard chemotherapy such as doxorubicin. We measured the IC-50 of combination of AMP-001/003 with doxorubicin in various triple negative breast cancer cells (MDA-MB-231, WHIM 12, BT 549). Under the experimental conditions, IC-50 for standard doxorubicin was around 388 nM. However, when the cells were incubated with AMP-001 or 002 (20 μM), there was a significant reduction of IC-50 for the combination from 388 nM level to 16-38 μM level (∼10 fold). This clearly indicates that AMP-001-002 enhance the potency of doxorubicin. This observation has a broader connotation that given the activation and inhibition of specific pathways by AMP derivatives (see mechanism of action), they evoke a better response from chemotherapy by reducing the eventual chemotherapeutic dose without compromising on efficacy, yet reducing the dose related toxicity. The change in IC-50 values was corroborated with photomicrographs of cell death (Fig 1, Supplementary).

It is recommended that multiple dose combinations of drugs have to be tested in order to map out synergistic or antagonistic effects and differentiation of additive Vs synergistic effects. The combination index is a standard method for two drug combinations to fix the ratio of doses at which synergicity occurs. The combination index was measured by generating isobologram for various doses of representative AMP-001 with doxorubicin or paclitaxel. The isobologram works by taking the effective concentration of each compound alone and drawing straight line between the axes. In this example (Fig 2B), the calculated ED90 for AMP-001 was ∼ 36 μM and for doxorubicin ∼2 μM and a straight line was drawn between these two pints. If the combination value is less than 1.00 and below the line would be considered as synergistic, while CI value above 1.00 and above the line would be considered as antagonistic in nature. The point on the line would be additive. The data clearly indicate at all the three ED50/75/90 values, the CI is <1.00 and below the line. This is quantitative data on the synergicity of AMP derivatives^29^.

Most tocopheryl derivatives are known to induce cell death through the generation of reactive oxygen species (ROS). Similarly, AMP-001-003 produced ROS, the kinetics of which followed a standard pattern raising the number of radicals initially (60-120 mins) followed by slow decline in about 6 hrs. However, cell death in cancer cells has to be correlated to downstream events such as release of apoptosis inducing factor (AIF-1) protein and Bax/Bcl2 expression. The biological targets involved for AMP-001/003 appear to be lysosomes and mitochondria. The uptake and distribution of acridine orange (AO) by cancer cells was assessed to determine the integrity of lysosome membrane permeabilization (LMP). In fact, acridine orange is a metachromatic lysosomotrophic agent that fluoresces red upon accumulation within the acidic environment of lysosomes and fluoresces green when located in the more alkaline cytosol or when bound to nuclear DNA. The induction of LMP leads to the release of acridine orange from the lysosomes out into the cytosol and nucleus. The reduction of red fluorescence upon incubation of AMP-001 or AMP-002 with PC3 or TNBC cells compared to control suggests that lysosome could be a potential target for ROS generated by AMP-001/003. It is significant that lysosomal membrane stability is related to the ability of ROS to overcome pro-survival autophagy^33^. The release of AIF-1 is yet another indication of irreversible cell death by AMP-001/003. Apoptosis inducing agents, in general act on mitochondria and provoke the permeabilization of the outer mitochondrial membrane, thereby triggering the release of potentially toxic mitochondrial proteins. Apoptosis-inducing factor (AIF), is a phylogenetically old flavoprotein which, in healthy cells, is confined to the mitochondrial intermembrane space^34^. Upon lethal signaling, AIF-1 translocates via the cytosol, to the nucleus where it binds to DNA and provokes caspase independent chromatin condensation. Mitochondrial and cytoplasmic cellular fractions of AMP-001/002 treated with TNBC MDA-MB-231 cancer cells are investigated using Western Blotting which indicates a significant decrease of AIF-1 in mitochondria at the expense of cytoplasmic AIF-1. α-TOS was used as a positive control. The release of apoptosis inducing factor-1, (AIF1) from mitochondria to cytosol is considered to be a non-reversible cell death signal correlated to CD95 cell death activation^35^.

Yet another useful downstream event of apoptosis is the upregulation of Bax and downregulation of Bcl2 protein. Apoptosis induction or resistance is reflected through Bax/Bcl2 ratio values (< 1, resistance, > 1 apoptosis). Taqman quantitative RT-PCR assay was used to estimate Bax/Bcl2 ratio with alpha-tocopherol as a negative control. The expression levels of GAPDH and Beta-Actin were used as normalizing house-keeping genes. The ratio of Bax/Bcl2 is > 1 (at 50 μM/24 hrs) for AMP-001 which indicates AMP-001 acts as an apoptotic enhancer facilitating the release of mitochondrial Cytochrome C into cytoplasm (See Supp 2), while the negative control did not show any effect. The effect is true with respect to both GAPDH and Beta-Actin controls. The observed significant increase of downstream Bax/Bcl2 ratio is well correlated to sensitization of cells which are <u>resistant to CD95</u> induced apoptosis^35^ which is an important correlation for establishing the potential mechanism of AMP-001/003.

### Potential Mechanism of Cancer Cells Sensitization

A) NF-kB Pathway: Cancer cells in particular, cancer stem cells (CSCs) and cancer resistant cells (CRCs) desensitize themselves to intervention by activating survival pathways (e.g. NF-kB) and downregulating cell death pathways (e.g. CD95). Conventional chemotherapy can kill bulk cancer cells leaving behind CSCs and CRCs which are responsible for recurrence of cancer. The initial step in acquiring the stemness properties by cancer cells is formation of mammospheres when cancers cells adhere together. To determine whether AMP-001 has an effect on self-renewal of CSCs, we performed mammosphere assay, a commonly used surrogate *in vitro* assay to measure self-renewal^36^. We used two cell line models. The first is the MCF-10A cells overexpressing SRC oncogene, which has previously been shown to possess cancer stem activity and susceptible to the effects of metformin^23^. The second is <u>mammary fat pad tumor-derived</u> MDA-MB-231 cells (TMD-231), which we had previously demonstrated to have CSC property and metastasizes spontaneously to lungs upon mammary fat pad injection^36^. The reduction of number of mammospheres was evident at 5 or 10 μM AMP-001 treatment. This effect of AMP-001 was similar to previously reported trans retinoic acid on stem cell activity of breast cancer cells including in TMD-231 cells^37^. NF-kB activation is correlated to the development of resistance to treatments. Hence, inhibition of NF-kB pathway selectively in cancer cells, potentially can make them susceptible to cell death. Conventional NF-kB inhibitors (curcumin, JSH 23) have similar activity at much higher μM range^38^ and are not targeted to CSCs leading to unwanted side effects.

### Potential Mechanism for Blocking Self-Renewal of CSCs

B). Inhibition of NF-kB Pathway: NF-kB is a transcription factor linked to the control of apoptosis signaling. Pharmacological inhibition of NF-kB is known to inhibit <u>proliferation of CSCs preferentially compared to bulk tumor</u> <u>population</u> suggesting that NF-kB activity is critical for the survival of CSCs^38^. Translocation of p65:p50 protein complex, representing canonical NF-kB, from cytoplasm to nucleus and its binding to the promoter regions of specific pro-survival genes linking epithelial to mesenchymal transition (EMT)/stemness phenotype is often observed in CSCs^40^. Overexpression of constitutively active p65 subunit of NF-kB in normal MCF-10A epithelial breast cells resulted in a dramatic shift to the CD44+/CD24-phenotype breast CSC “stemness” behavior linked to mammosphere forming ability^37^. We have assessed NF-kB inhibition using Western Blotting through nuclear and cytosolic extracts from TNBC cells treated with AMP-001/002. Quantitative data in Fig 7D shows a significant <u>decrease in nuclear p65</u> fractions with concomitant <u>increase in</u> <u>cytoplasmic</u> p65 fractions upon AMP-001/002 treatment compared to untreated control (Fig 7D, red line 1 Vs 3-4) or positive control α-tocopherol succinate TOS (lane 2). This clearly indicates NF-kB pathway may be one of the targets of AMP-001/002, which could partially explain the effects of the drugs on CSCs (Fig 6). Deliberate NF-kB activation by alfa-TNF was nullified by AMP-001/003 indicating the potency of AMP-001/003 to inhibit NF-kB pathway. This result is very important since NF-kB activation is the survival pathway for CSCs and the controlled NF-kB inhibition is known to sensitize cells to chemotherapy^38^.

Sensitizing tumor cells to chemotherapy has an additional advantage that chemotherapy dose can be tuned down at the expense of sensitizing agent to yield a new formulation which will yield low toxicity due to low dose. The downregulation of cell death pathway CD95 is correlated to the development of resistance. Fas resistant MDA-MB-231 TNBC cells were treated with different concentrations of AMP-001 to assess the expression of CD95 (43 KDa) by Western blotting (Fig. 8). The significant change in the intensity of the CD95 band in the isolated membrane fractions of treated cells indicates a possible translocation of CD95 receptor from cytosol to the surface of cell which triggers the apoptotic signal. In contrast, the untreated cells showed low intensity CD95 band. Translocation of CD95 from cytosol to the membrane is a common observation for CD95 Trail synergy^31^ which I further correlated to NF-kB inhibition (e.g. which influences the cellular sensitivity to the Fas death receptor pathway). Activation of CD95 pathway is important particularly, for CRCs and for CRCs where significant down regulation of CD95 is reported.

Improving the clinical performance of the current treatments for cancer in general and TNBC in particular is the main objective for sensitizing low-responsive tumor cells. Doxorubicin is one of the front-line treatments for TNBC patients. Doxorubicin treatment has been implicated in the hyperactivation of PARP in heart leading to cardiotoxiciy^41^. The inhibition of PARP by PARP inhibitors (e.g. Abbott ABT472) is known to sensitize tumor cells which are otherwise nonresponsive to doxorubicin, potentiated the effect of doxorubicin for tumor cells and ameliorated cardiotoxicity^42^. Dose dependent incubation of AMP-001 with MDA-MB-231 cells and analysis by Western blotting (Fig 8B) showed a clear cleavage of PARP at 89 KDa which suggests that PARP function may be inhibited which is correlated to the activation of pro-apoptotic caspases and subsequent apoptosis. This may also be related to sensitization of tumor cells by AMP-001.

The relative toxicity profiles of AMP-001/003 with respect to doxorubicin, particularly in cardiomyocytes and normal breast cells would help us to optimize the dose regimen in tumor animal models. Hence, induced pluripotent stem cell derived human cardiomyocytes (iPSC, Ionic Transport Assays, LLC) were treated with AMP-001/002 *in vitro* which showed high IC-50 value (> 200 μM) for AMP-001/002, while doxorubicin is cardiotoxic at as low dose as at 9.6 μM (Fig 9). At the same time, IC-50 values for AMP-001/002 are in 20-40 μM range for cancer cells (Table 1). The high differential IC-50 of AMP-001 between TNBC cells Vs cardiomyocytes (32 μM Vs > 250 μM respectively) may help to establish high target/non-target potential *in vivo*. It is to be noted that standard oncology drugs Sorafinib or doxorubicin are cardiotoxic at early μM range compared to AMP-001/002.

Efficacy and toxicity are <u>the</u> two main criteria for a successful clinical translation. Six-week-old female Nu/Nu rats (n = 8) were innoculated in the mammary fat pad with 5×10^6^ MDA-MB-231 cells stably expressing firefly luciferase. Tumors were allowed to reach 100-150 mm^3^ before i.p administration of AMP-001. For *in vivo* studies, six-week-old female Nu/Nu mice (n = 8) were innoculated in the mammary fat pad with 5×10^6^ MDA-MB-231 cells stably expressing firefly luciferase. The treatment schedule for this experiment was AMP-001 monotherapy at 100 mg/Kg or 200 mg/Kg dose x 3 days x 2 weeks. Tumors were monitored by bioluminescence (IVIS 200, Perkin Elmer). No toxicity was detected as far as significant weight loss, decreased mobility, grooming behavior or labored breathing are concerned. Fig. 10 A shows a significant reduction in the bioluminescent signal correlated to tumor volume plotted as a percentage of tumor growth normalized at day 1 of injection (Fig. 10B, p < 0.05) compared to untreated control. The classic V curve confirms tumor regression “*In vivo*” and little adverse effect on heart (Fig 10C). Data are similar on kidney, liver and spleen sections (not shown). It is to be noted that low dose combination of AMP-001/002 (∼10-20 μM) is synergistic with doxorubicin *in vitro*, while high dose AMP-001 yielded tumor regression on its own with low off-target toxicity. Although, the final therapeutic dose 200mg/Kg seems to be high to make it commercially successful, the efficacy with no significant toxicity to non-target organs is a significant progress in the management of cancer disease. Also, AMP-001/003 are activating and inhibiting specific pathways to sensitize tumor cells, it is proposed to use them as a selective tumor sensitizer and hence, low dose of AMP-001/003 becomes a part of the combination formulation. Further studies on determination of maximum tolerable dose (MTD) which determines therapeutic index, leverage for tuning dose regime, extensive pharmacokinetic and pharmacodynamic aspects of drug development are in progress.

## Conclusions

Since cancer is a group of diseases, several targets and pathways are involved in the progression of the disease. The combination of AAAPT leading molecule with a conventional chemotherapy brings multiple pathways involved in cancer cells which may help offset several dysregulated pathways for a potential synergicity.

In summary, the effects of novel vitamin E analogue AMP-001 is as a preclinical candidate is described. AMP-001 enhanced doxorubicin induced apoptosis in several cancer cells including TNBC cells via ROS-dependent mitochondrial apoptotic pathway, activation of cell death pathway (CD95) and inhibition of survival pathways (NF-kB and PARP). The efficacy of doxorubicin antitumor activity can be amplified by combining it with AMP-001, which would permit lower dose use in patients without affecting the positive clinical outcome. These effects are confirmed both *in vitro* and *in vivo*. Our current studies indicated that AMP-001 might be a potential candidate in synergetic with the existing FDA approved chemotherapeutic drugs for treatment of TNBC cancer which has limited number of treatments. Ultimately, the combination treatments success depends on how much low dose can be made effective in a clinical situation, reduced overall toxicity and improved patient compliance. Extending the synergicity of AMP-001 to other chemotherapeutics (e.g. Gemcitabine, Paclitaxel, Carboplatin, Avastin etc.) and other treatment modalities such as radiation, radionuclide therapy and immunotherapy may be significant contribution to cancer research. Further studies are planned to optimize the dose combination as to which one has higher efficacy and low toxicity compared to the existing treatments.

## Acknowledgements

Part of this research was supported by the grant SBIR NIH R43CA183353.

## Conflict of interest

The authors declare no competing financial interest.

